# Inhibition of neutrophil degranulation by Nexinhib20 delays the development of radiation-induced pulmonary fibrosis

**DOI:** 10.1101/2025.10.22.684057

**Authors:** Stefanie Ruhland, Konstantina Nikolatou, Victoria L. Bridgeman, Tatiana Rizou, Erik Sahai, Jayanta Bordoloi, Emma Nolan, Alberto Elosegui-Artola, Ilaria Malanchi

## Abstract

Despite advances in radiation delivery techniques that enhance tumour targeting and minimise collateral exposure, healthy tissues remain susceptible to radiation-induced fibrosis, a chronic and progressive condition that can severely compromise patient quality of life. Therapeutic options to prevent or treat radiation-induced fibrosis remain limited. Recent work has shown that neutrophils infiltrating healthy lung tissue after irradiation adopt an activated phenotype capable of perturbing cellular responses of both epithelial and mesenchymal cells. However, the contribution of these radiation-educated neutrophils to the development of radiation-induced fibrosis remains unclear. Using targeted, image-guided lung irradiation to deliver a dose sufficient to induce fibrosis within four months, we demonstrate that neutrophils are essential for the efficient development of clinically evident fibrosis. Lung irradiation educates neutrophils, enabling them to promote early alterations in the extracellular matrix shortly after radiation exposure. We further show that this “educated” phenotype depends on neutrophil degranulation activity. Pharmacological inhibition of degranulation with Nexinhib20 redirected these cells toward a pro-angiogenic and anti-fibrotic phenotype. Importantly, this treatment was also associated with a transcriptional shift in mesenchymal cells away from a pro-fibrotic program, resulting in a marked delay in the onset of radiation-induced fibrosis. Importantly, Nexinhib20 did not impair the efficacy of cancer radiotherapy, underscoring its potential as a therapeutic strategy to prevent fibrotic complications without diminishing anti-tumour effectiveness.

## Introduction

Radiotherapy is a cornerstone of cancer treatment, with approximately half of all cancer patients receiving radiotherapy during the course of their disease (1). As ionising radiation affects both malignant and normal cells, treatment schedules and dosing must balance tumour radiosensitivity with healthy tissue toxicity. Despite advances in delivery techniques that improve tumour targeting and reduce collateral exposure, healthy tissues remain vulnerable with toxicities depending on radiation dose, fractionation, irradiated volume and tissue-specific radiosensitivity (2–4). The lung, one of the most radiosensitive organs, is particularly susceptible, with 16–28% of patients with thoracic malignancies developing radiation-induced pulmonary fibrosis following radiotherapy (2). This progressive and irreversible condition is characterised by scarring, chronic respiratory symptoms, reduced lung function and substantially impaired quality of life (2). With improving cancer survival rates, the absence of effective treatment or preventative strategies makes radiation-induced fibrosis an increasingly urgent clinical challenge.

Radiation, a sterile injury, damages lung tissue and perturbs tissue integrity. Such a tissue injury normally triggers a tightly regulated repair response, restoring structure and function through coordinated activation of epithelial, mesenchymal and immune cells. When repair mechanisms become imbalanced, pro-fibrotic pathways are activated, stimulating fibroblast proliferation and differentiation into myofibroblasts, leading to excessive extracellular matrix (ECM) deposition, fibre cross-linking and matrix stiffening (5–9). These changes disrupt normal lung architecture and, over time, drive irreversible scarring. Foamy (lipid-laden) macrophages emerge from altered lipid uptake and further support fibroblast activation and ECM remodelling, exacerbating fibrosis (10–13). Mice deficient in the lipid uptake receptor CD36 are partially protected from pulmonary fibrosis in bleomycin models, highlighting their pathological relevance (14).

Neutrophils are key mediators of tissue repair, clearing debris, scavenging pro-inflammatory signals and releasing pro-resolving factors to support re-epithelialisation and tissue regeneration (15–21). For example, in sterile thermal hepatic injury, neutrophils actively contribute to regeneration by clearing damaged vessels and promoting revascularisation (18) . Similarly, they facilitate cardiac repair after myocardial infarction (22) and modulate repair kinetics in ventilator-induced lung injury, where neutropenia delays tissue recovery (23) . Together, these studies underscore the essential contribution of neutrophils to the resolution of injury. Effective resolution requires that their activity is tightly regulated, for instance through their removal by macrophages or reverse migration, which reinforces an anti-inflammatory and regenerative environment (18,19,24–26). However, excessive or prolonged neutrophil activation, including degranulation and reactive oxygen species (ROS) release, can amplify tissue injury and switch their beneficial functions towards tissue damage (19,20,27,28).

We previously demonstrated that neutrophils infiltrating healthy lung tissue after irradiation adopt an activated “radiation-educated” phenotype, capable of altering the tissue microenvironment and impacting both epithelial and mesenchymal cells (29). Given this broad impact of neutrophils on early post-radiation tissue responses, in this study we examined whether radiation-educated neutrophils may also contribute to the development of radiation-induced fibrosis.

## Results

### Neutrophils support the development of radiation-induced fibrosis

To evaluate the contribution of neutrophils to the development of radiation-induced pulmonary fibrosis, we examined fibrosis progression in genetically neutropenic FVB granulocyte colony stimulating factor (GCSF) knockout (ko) mice, in which neutrophil-mediated injury responses are impaired and pulmonary neutrophil presence is markedly reduced (Fig. 1A and B). Using CT-guided targeted irradiation, 13 Gy was delivered to the right lung (targeted irradiated lung, IR), while the left lung (off-target, offIR) was largely spared, with only minimal scatter exposure (10% of the tissue receiving ≤1 Gy) (Supp. Fig. 1A and B). Fifteen weeks post-irradiation, Picrosirius Red staining revealed pronounced fibrosis in the targeted irradiated lungs, characterised by extensive collagen deposition and accumulation of foam cells, aberrant CD68^+^ macrophages expressing the lipid scavenger receptor CD36 (Fig. 1C-E and Supp. Fig. 1C and D). These foamy macrophages are reported to support the development of fibrosis by secreting profibrotic mediators and stimulating fibroblast activation (10–12,14). Those fibrotic changes and foamy macrophages were restricted to the targeted irradiated lung tissue (Fig. 1E).

**Figure 1:**
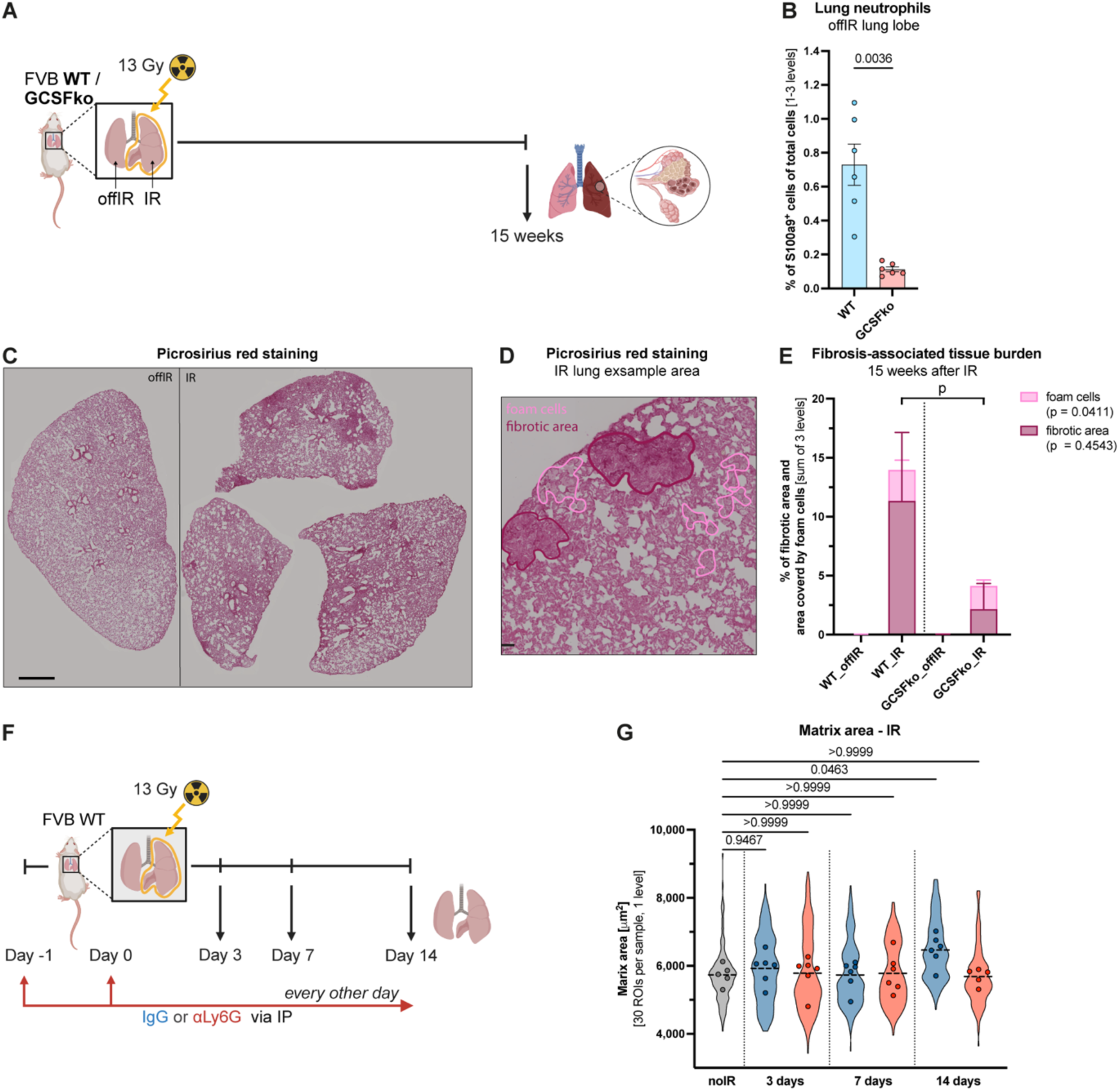
Neutrophils support the development of radiation-induced fibrosis. **(A)** Schematic illustrating the experimental setup used to evaluate the role of neutrophils in the development of radiation-induced pulmonary fibrosis. **(B)** Quantification of neutrophil presence in offIR lung tissue by counting S100a9^+^ stained cells as a percentage of total cells across slides from one to three levels. Statistical analysis was performed using Welch’s t-test. Data are presented with mean ± SEM (n = 6). **(C)** Representative image of picrosirius red stained lung tissue slides showing the non-targeted left offIR lung and the targeted right IR lung. Scale bar: 1000 µm. **(D)** Representative ROI of IR lung stained with picrosirius red, with fibrotic regions and areas covered by foamy macrophages outlined. Example of area measurements. Scale bar: 50 µm. **(E)** Quantification of fibrotic area and area covered by foamy macrophages. Values represent the sum over three levels divided by the total lung area. Statistical comparison of foam cells or fibrotic area between WT_IR and GCSFko_IR was calculated on graphs in Supp. Fig. 1E and F. Data shown as median + IQR. **(F)** Schematic illustrating the experimental setup used to evaluate the role of neutrophils on early matrix changes after radiation exposure. **(G)** Quantification of matrix area across 30 ROIs (48,163 µm²) from one level per sample using TWOMBLI. Violin plots show the distribution of ROIs and dots indicate the average value for each mouse. Statistical analysis was performed on individual mice using one-way ANOVA against non-irradiated (noIR) control samples. Data are presented with mean (n = 5-6).

Strikingly, GCSFko mice, despite an unchanged abundance of foamy macrophages, showed a marked reduction in fibrotic area (Fig. 1E and Supp. Fig. 1E and F). Automated TWOMBLI analysis, which quantifies matrix area within regions of interest (ROIs) (Supp. Fig. 1G), confirmed these findings, detecting increased matrix area in response to radiation, with significant expansion observed only in wildtype (WT) mice (Supp. Fig. 1H). Together, these results suggest that neutrophils are key drivers of radiation-induced fibrosis.

Neutrophil infiltration peaks in the lung seven days post-exposure (29), therefore, we next investigated whether neutrophils also contribute to early ECM changes that may set the stage for long-term fibrosis development. To directly test the effect of irradiation on the ECM in absence of neutrophils, we used an αLy6G antibody (Supp. Fig. 2A), to efficiently deplete neutrophils for a short period without the broader alterations associated with the GCSFko genetic model. The depletion was initiated one day prior to irradiation and remained effective over the seven-day period, during which increased neutrophil infiltration was observed at day seven (Supp. Fig. 2B), coinciding with an immune cell recruiting and inflammatory signature of the mesenchymal population (Supp. Fig. 2C).

Matrix analysis over time revealed an immediate, neutrophil-independent ECM response, reflected by a transient increase in matrix area, likely a result of matrix contraction, that subsided within 24 h (Supp. Fig. 2D-G). This early response was not restricted to the targeted irradiated region of the lung and is therefore unlikely to be relevant to fibrosis development (Supp. Fig. 2G). Whilst no striking changes were observed up to one week after radiation exposure, two weeks after a significant difference in the matrix expansion was observed in the control mice but absent in the neutrophil-depleted group (Fig. 1F and G and Supp. Fig. 2H and I). Treatment with the neutrophil depletion antibody effectively reduced neutrophil presence for up to two weeks (Supp. Fig.2J), with no effects observed in the off-target regions or due to the antibody treatment itself (Supp. Fig.2I). Collectively, these results indicate that neutrophils contribute to radiation-induced fibrosis from early time points by influencing initial ECM remodelling, which ultimately drives fibrotic development.

### Radiation-educated neutrophils are phenotypically altered

Since neutrophils contribute to radiation-driven ECM changes yet are not directly exposed to the radiation themselves (29), we next investigated whether neutrophils isolated from pre-irradiated lungs, referred to as radiation-educated, display distinct phenotypic features compared to those from non-irradiated (noIR) control lungs. First, we analysed surface marker expression by flow cytometry on neutrophils isolated one or two weeks after irradiation, with their abundance peaking at day seven and returning towards baseline levels by day 14 (Fig. 2A and 2B). Radiation-educated neutrophils progressively upregulated CD11b expression, indicative of activation, displayed a transient reduction in the maturation marker CD101 at day seven and exhibited increased CXCR4 levels by day 14 (Fig. 2C-E), suggesting persistence of an altered phenotype and potentially increased aged status.

**Figure 2:**
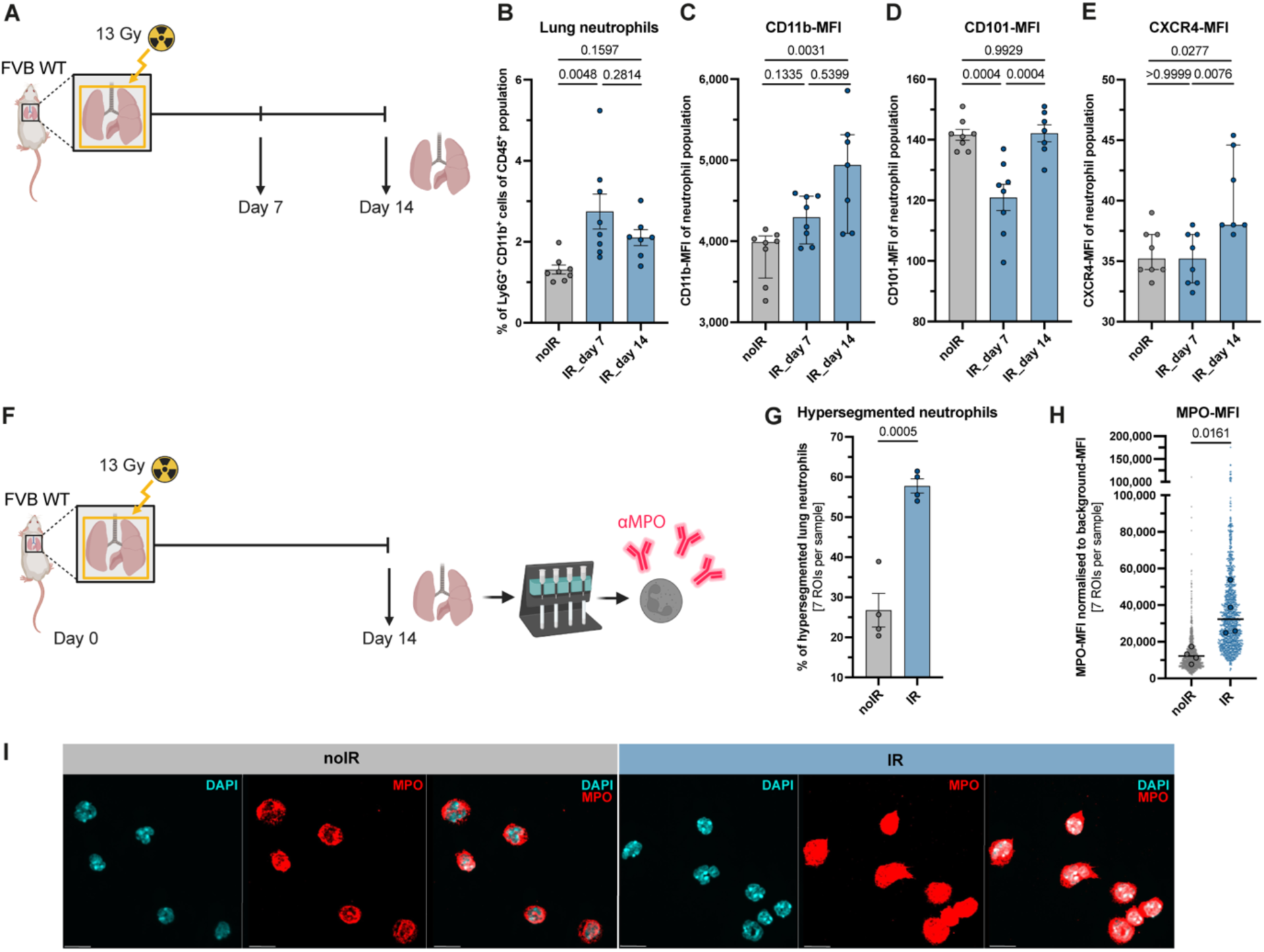
Radiation-educated neutrophils display altered phenotype. **(A)** Schematic illustrating the experimental setup used to evaluate changes of neutrophil surface marker expression at seven and 14 days after irradiation by FACS. **(B-E)** Percentage of neutrophils among CD45⁺ immune cells in lung tissue (B) and their expression levels of surface markers CD11b (C), CD101 (D) and CXCR4 (E), quantified by FACS. Statistical analysis was performed using one-way ANOVA for normally distributed data (B and D) or Kruskal-Wallis test for non-normally distributed data (C and E). Data are presented with mean ± SEM (B and D) or median ± IQR (C and E) (n = 7-8). **(F)** Schematic illustrating the experimental setup used to evaluate changes in nuclear segmentation and MPO expression levels of neutrophils isolated from pre-irradiated lung tissue 14 days afterwards by immunofluorescence. **(G)** Quantification of hypersegmented neutrophils from 7 ROIs per biological replicate. Neutrophils were classified as hypersegmented if the nucleus contained ≥6 lobes. Statistical analysis was performed using an unpaired t-test. Data are presented with mean ± SEM (n = 4). **(H)** Quantification of MPO-MFI per neutrophil across 7 ROIs per biological replicate (same neutrophils as in G). Each neutrophil is shown as a small dot in the background (forming a violin plot), with the average value per mouse shown as a larger dot in front. Statistical analysis was performed on the per-mouse averages using an unpaired t-test. Data are presented with mean (n = 4). **(I)** Representative images of neutrophils isolated from noIR or IR lung tissue 14 days after irradiation, stained with DAPI and MPO. Scale bar: 10 µm.

Second, immunofluorescence analysis of neutrophils 14 days post-irradiation revealed a shift toward hypersegmented nuclear morphology and elevated myeloperoxidase (MPO) content compared to controls (Fig. 2F-I). The increased MPO levels suggests that radiation-educated neutrophils have an enhanced potential to modify the ECM, consistent with their contribution in early matrix remodelling. Taken together, radiation-educated neutrophils exhibit phenotypic changes indicative of elevated activation and granule protein enrichment post-irradiation, consistent with their contribution to ECM remodelling and subsequent fibrosis development.

### Neutrophil degranulation inhibition with Nexinhib20 prevents early matrix changes

Since radiation-educated neutrophils exhibited increased granule protein content, we next asked whether the absence of individual granule proteins would be sufficient to prevent early matrix changes. We focused on neutrophil elastase (NE) and MPO, two granule proteins with the potential to influence ECM integrity and remodelling (30,31). MPO was strongly upregulated in our immunofluorescence analysis (Fig. 2H) and was also previously identified as one of the most markedly elevated proteins in radiation-educated lung neutrophils, showing a 2-fold elevation relative to their non-irradiated controls (29). While NE displayed only a modest enhancement (1.12-fold) in the same study, previous work has reported increased NE levels in bronchioalveolar lavage fluid from patients with pulmonary fibrosis (32,33). We therefore assessed the functional contribution of MPO and NE by applying our irradiation protocol to mice lacking MPO (MPOko) or functional NE (Ela2ko) respectively and compared them with WT controls (Fig. 3A). Differences in genetic background did not affect baseline matrix area, as no changes were observed in the off-target lung samples (Supp. Fig. 3A). In WT and Ela2ko mice, irradiated lungs consistently displayed increased matrix area relative to their paired off-target lungs (Fig. 3B and C). In contrast, MPOko mice showed a heterogeneous response (Fig. 3D). While some animals exhibited a pronounced increase in matrix area, others showed only modest elevation or even a reduction, resulting in no significant difference overall. Due to the variable response of MPOko mice and the persistent matrix increase in Ela2ko mice, these results showed that the loss of a single granule protein was not sufficient to prevent early ECM remodelling, as the matrix areas of their irradiated lungs were comparable to those of WT animals (Supp. Fig. 3B).

**Figure 3:**
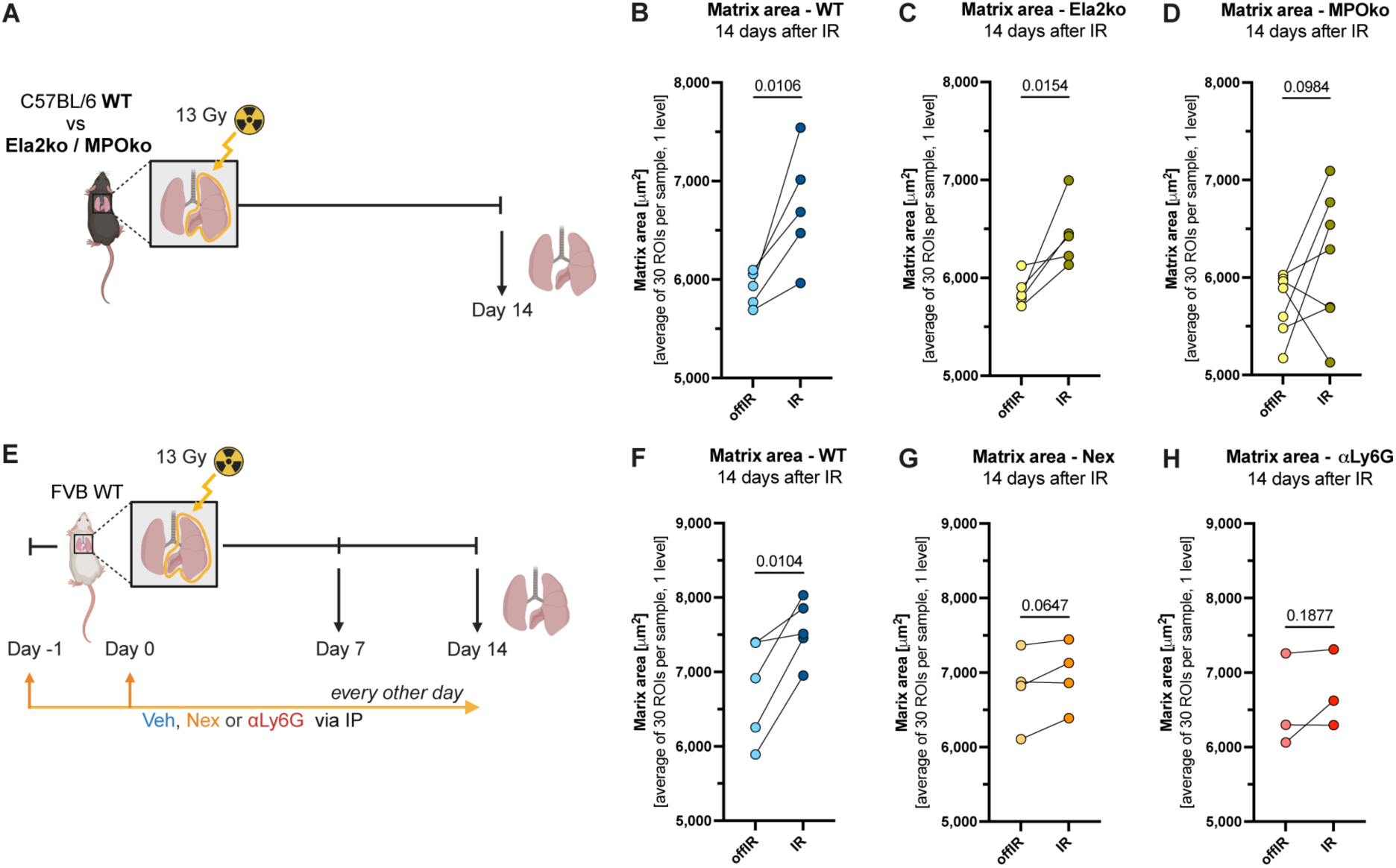
Blocking degranulation with Nexinhib20 prevents early matrix changes. **(A)** Schematic illustrating the experimental setup used to evaluate early matrix changes in genetically modified mice with altered granule composition. **(B-D)** Quantification of matrix area across 30 ROIs (48,163 µm²) from one level per sample using TWOMBLI. Dots show the average per mouse, with paired offIR and IR lungs indicated. Statistical analysis was performed using paired t-test (n = 5-7). **(E)** Schematic illustrating the experimental setup used to evaluate the effect of the degranulation inhibitor Nexinhib20 on early matrix changes. **(F-H)** Quantification of matrix area across 30 ROIs (48,163 µm²) from one level per sample using TWOMBLI. Dots show the average per mouse, with paired offIR and IR lungs indicated. Statistical analysis was performed using paired t-test (n = 3-5).

We next decided to broadly inhibit granule protein release by using the degranulation inhibitor Nexinhib20. This compound blocks the interaction between Rab27a and JFC1, thereby preventing the degranulation of azurophilic granules (34), containing MPO and NE among others (Supp. Fig. 3C). This approach enabled simultaneous targeting of multiple granule proteins, thereby preventing their interaction with the ECM. Supporting the relevance of this strategy, Rab27a expression showed a 1.4-fold increase in radiation-educated neutrophils seven days post-irradiation compared to controls (Supp. Fig. 3C, proteomics data from Nolan et al. (2022) (29)), suggesting that Nexinhib20 treatment could mitigate early ECM changes.

The short-term changes in ECM were assessed when targeted irradiation was administered in presence of Nexinhib20 treatment, vehicle control (Veh) or neutrophil-depleting antibody αLy6G (Fig. 3E and Supp. Fig. 3D and E). In accordance with our previous experiment (Fig. 1G), no consistent irradiation-induced changes in matrix area were observed one week after radiation (Supp. Fig. 3F-H), whereas by two weeks, Veh-treated mice showed a significant, uniform and robust increase in matrix area (Fig. 3F). Importantly, this increase was absent in both Nexinhib20- and αLy6G-treated mice (Fig. 3G and H). While direct comparison of irradiated lungs across the three groups revealed no differences at the seven day timepoint, two weeks after irradiation, Veh-treated mice displayed a clear trend towards higher matrix area (Supp. Fig. 3I and 3J). Of note, Nexinhib20 treatment itself did not affect matrix area, as off-target lungs were comparable between groups at both time points (Supp. Fig.3K and L).

### Nexinhib20 modulates the radiation-educated neutrophil phenotype and transcriptomic signature

Given that Nexinhib20 effectively prevented early matrix changes, we next asked whether it also modulates the phenotype of the radiation-educated neutrophils, which we previously found to exhibit increased activation, aging markers as well as higher proportion of hypersegmented neutrophils (Fig. 2). While Nexinhib20 treatment did not affect the radiation-driven elevated surface expression of CD11b, it prevented the decrease in CD101 surface expression at day seven, with no change at day 14 (Supp. Fig. 4A-D). Interestingly, CXCR4 surface expression showed a trend towards downregulation at day 14 in the context of Nexinhib20 treatment (Supp. Fig. 4E and F). Assessment of hypersegmentation via immunofluorescence 14 days after irradiation revealed that Nexinhib20 also averted the irradiation-induced increase in hypersegmented neutrophils, maintaining the levels comparable to non-irradiated controls (Fig. 4A-C). To assess functional consequences, neutrophil motility was quantified by plating cells on a gelatin matrix and tracking their movement via time-lapse confocal microscopy (Fig. 4D). Radiation-educated neutrophils from Veh-treated mice exhibited increased motility, whereas this effect was strongly attenuated in neutrophils from Nexinhib20-treated mice (Fig. 4E and F). Together, these results indicate that Nexinhib20 treatment mitigates some of the phenotypic changes that typically occur in a pre-irradiated environment.

**Figure 4:**
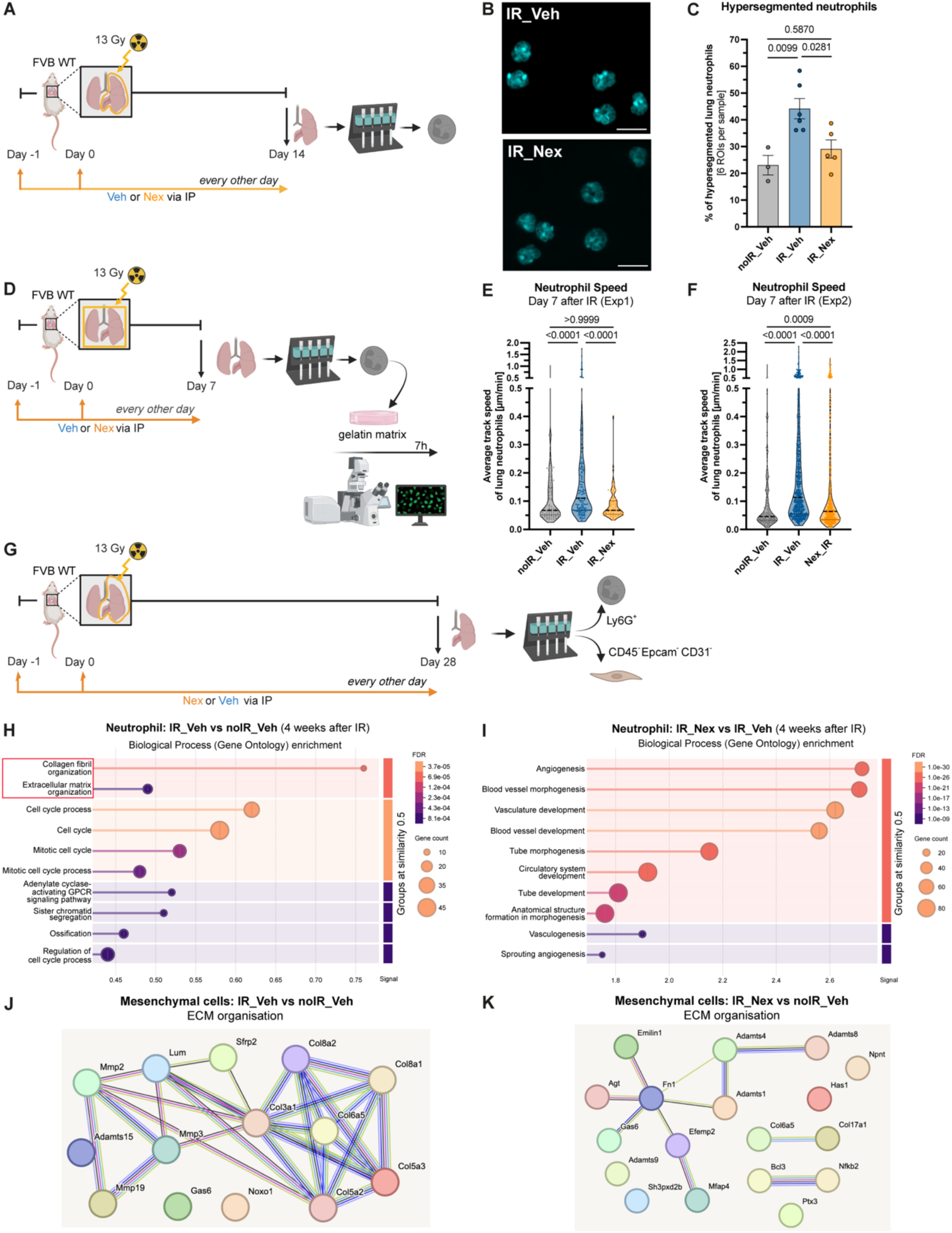
Nexinhib20 modulates the radiation-educated neutrophil phenotype and transcriptomic signature. **(A)** Schematic illustrating the experimental setup used to evaluate changes in nuclear segmentation of neutrophils isolated from pre-irradiated lung tissue 14 days afterwards by IF. **(B)** Representative images of neutrophils isolated from Veh-treated or Nex-treated lung tissue 14 days after irradiation, stained with DAPI. Scale bar: 10 µm. **(C)** Quantification of hypersegmented neutrophils from six ROIs per biological replicate. Neutrophils were classified as hypersegmented if the nucleus contained ≥6 lobes. Statistical analysis was performed using an unpaired t-test. Data are presented with mean ± SEM (n = 3-6). **(D)** Schematic illustrating the experimental setup used to evaluate changes in the average track speed of neutrophils isolated from pre-irradiated lung tissue seven days afterwards and placed on a gelatin matrix by time-lapse confocal microscopy. **(E and F)** Quantification of the average neutrophil track speed over a timeframe of 7h. In experiment one neutrophils were collected from one biological replicate and in experiment two neutrophils were collected from three biological replicates with the violin showing the distribution and each neutrophil indicated by a dot. Statistical analysis was performed using a Kruskal-Wallis test. Data are presented with median ± IQR. **(G)** Schematic illustrating the experimental setup used to isolate neutrophils and mesenchymal cells four weeks post-irradiation for bulk RNAseq. **(H and I)** GO-term analysis of differential expressed gene analysis of bulk RNAseq of neutrophil isolated from Veh- (G) or Nex-treated IR (H) vs Veh-treated noIR lung tissue four weeks post-irradiation via the STRING pathway analysis webpage (35). **(J and K)** Enriched signature of ECM organisation gene set within the mesenchymal cell population either enhanced in Veh-treated (J) or Nexinhib20-treated (K) IR sample over noIR control sample (generated with (35)).

We next investigated whether differences are also reflected in their transcriptional programs. Bulk RNAseq was performed four weeks post-irradiation to capture sustained gene expression changes in both neutrophils and the mesenchymal compartment (Fig. 4G). Both populations were isolated from the same lung samples for the irradiated groups, by first positive selecting for Ly6G^+^ neutrophils via Magnetic activated cell sorting (MACS) and then collecting the mesenchymal cells as the CD45^-^ Epcam^-^ and CD31^-^ fraction, which is largely composed of fibroblasts, key regulators of ECM production and maintenance. Those samples were compared to neutrophils and mesenchymal cells from non-irradiated Veh controls lungs, which were obtained from separate mice.

In non-irradiated lungs, neutrophils displayed a strong inflammatory gene signature (Supp. Fig. 4G). Radiation-educated neutrophils, however, showed reduced inflammatory gene expression and increased cell cycle related genes (Supp. Fig. 4G). The ECM organisation gene signature was upregulated in radiation-educated neutrophils compared to control lung neutrophils (Fig. 4H and Supp. Fig. 4G). Radiation-educated neutrophils isolated from Nexinhib20-treated mice also exhibited enrichment of ECM organisation–related genes (Supp. Fig. 4G). However, compared with radiation-educated neutrophils from Veh-treated animals, they displayed a transcriptional shift toward a vascular development signature (Fig. 4I and Supp. Fig. 4G). Thus, four weeks post-irradiation, radiation-educated neutrophils adopt an anti-inflammatory and ECM remodelling transcriptional profile, while Nexinhib20 shifts them toward a more pro-angiogenic state, extending beyond ECM-related pathways.

Analysis of the mesenchymal compartment revealed four clusters, two of which expressed either erythrocyte-specific genes (Hba-a, Alas2, Slc4a1) or epithelial markers (EpCAM, Sftpc, Abca3, Napsa), likely reflecting residual contamination, particularly in non-irradiated controls (Supp. Fig. 4H). The remaining clusters were characterised by inflammatory or ECM organisation gene signatures, with both signatures being low in non-irradiated controls (Supp. Fig. 4H). As expected, lung irradiation induced a strong ECM organisation program in mesenchymal cells, characterised by collagen production and matrix-modifying enzyme release (Fig. 4J and Supp. Fig. 4H and I). In contrast, Nexinhib20 treatment shifted mesenchymal cells toward an immune-responsive and immune-regulatory state, while reducing the ECM organisation signature (Supp. Fig. 4H and J). Interestingly, residual ECM-related genes were enriched for fibronectin, elastic fibre and proteoglycan-processing pathways, emphasising structural integrity, vascular scaffolding and matrix elasticity rather than bulk collagen production (Fig. 4K).

Collectively, Nexinhib20 mitigates the radiation-educated neutrophil characteristics and shifts their transcriptional program from one primarily focused on ECM organisation to a strongly pro-angiogenic profile. This shift is mirrored in the mesenchymal compartment, where fibroblasts transition from a pro-fibrotic phenotype characterised by bulk collagen production and matrix-remodelling enzyme release to an immune-responsive state focused on maintaining matrix elasticity and supporting vascular scaffolding. This reflects a transition from matrix stiffness toward structural integrity.

### Nexinhib20 delays the development of radiation-induced fibrosis and enhances cancer radiotherapy efficacy

Building on the Nexinhib20-driven long-term transcriptional shifts observed in radiation- educated neutrophils and in the mesenchymal compartment, which collectively promote a less pro-fibrotic phenotype, we next investigated whether these changes translate into reduced development of clinical fibrosis. Using our established radiation model, mice were treating with Nexinhib20 or its Veh control for eight weeks (Fig. 5A). By this time point, fibrotic areas and foamy macrophages first become detectable, marking the onset of clinical fibrosis (Supp. Fig. 5A and B). A group of GCSFko mice receiving Veh was included to compare the effect of reduced neutrophil numbers with the degranulation inhibitor (Fig. 5A). At eight weeks, fibrotic area was minimal and comparable between groups (Supp. Fig. 5A). However, both Nexinhib20-treated and GCSFko mice showed a trend towards reduced foamy macrophage abundance relative to the Veh-treated WT mice (Supp. Fig. 5B). As foamy macrophages promote local fibrosis progression (10–12,14), this reduction may indicate extenuated fibrosis development in these groups.

**Figure 5:**
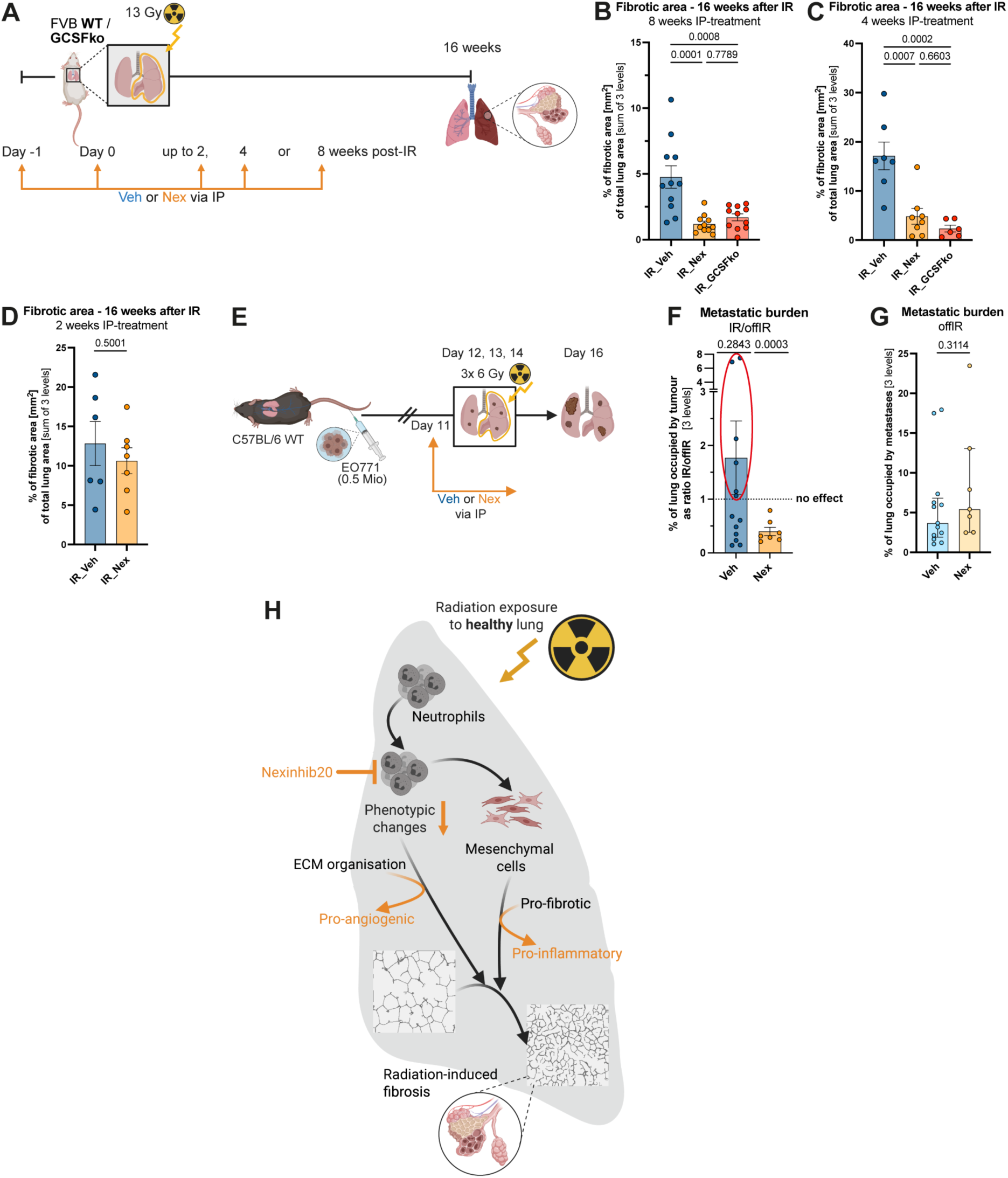
Nexinhib20 as a potential therapeutic strategy to delay radiation-induced fibrosis without compromising cancer radiotherapy efficacy. **(A)** Schematic illustrating the experimental setup used to evaluate the effect of Nexinhib20 on the development of radiation-induced pulmonary fibrosis with treatment periods of two, four or eight weeks. **(B-D)** Bar charts showing individual mouse data for the sum of fibrotic area in the IR lung over three levels. Mice received Nexinhib20 or Veh for either eight weeks (B), four weeks (C) or two weeks (E). All lungs were harvested 16 weeks post-irradiation. Statistical analysis was performed using one-way ANOVA (B and C) or unpaired t-test **(E).** Data are presented with mean ± SEM (n = 6-11). (E) Schematic illustrating the experimental setup used to evaluate the effect of Nexinhib20 on the efficiency of cancer radiotherapy in the setting of experimental EO771 breast cancer metastases in the lung. **(F)** Quantification of the effect of radiotherapy on the metastatic burden as ratio of the burden of targeted IR vs non-targeted offIR lung side with one indicating no change in the metastatic burden between right and left lung side. Statistical analysis was performed using a one sample t and Wilcoxon test against one. Data are presented with mean ± SEM (n = 7-11). **(G)** Comparison of metastatic burden between Veh- and Nex-treated non-targeted offIR lungs. Statistical analysis was performed using Mann-Whitney test. Data are presented with median ± IQR (n = 7-11). **(H)** Proposed model of neutrophils supporting the development of radiation induced fibrosis. Post-irradiation neutrophils acquire a radiation-educated phenotype that promotes fibrotic progression. Treatment with the neutrophil degranulation inhibitor Nexinhib20 delays fibrosis by diminishing the phenotypic changes and switching the neutrophil signature toward a pro-angiogenic profile, which is accompanied by a shift of mesenchymal cells from a pro-fibrotic to a pro-inflammatory state, reducing extracellular matrix accumulation and enhancing matrix integrity.

To test this conclusion, mice were again treated for eight weeks with lungs now being collected 16 weeks post-irradiation, when fibrosis is well established (Fig. 5A). At this stage, Nexinhib20- treated mice showed a significant decrease in fibrotic area compared to Veh-treated WT mice, reaching levels similar to those observed in GCSFko animals (Fig. 5B). Shortening Nexinhib20 treatment to four weeks (Fig. 5A) revealed that this regimen retained full protective efficacy, similar to GCSFko animals (Fig. 5C). However, a two-week treatment proved insufficient, thereby defining the clinically relevant time window required for effective fibrosis prevention (Fig. 5D). No changes in the abundance of foamy macrophages were observed across these experiments (Supp. Fig. 5C-E). These results align with the time point of our bulk RNAseq analysis, which showed that neutrophils shifted from a pro-fibrotic to a pro-angiogenic transcriptional profile after four weeks of treatment (Fig. 4G–I). This shift appears to establish a stable, less pathological neutrophil state that no longer supports the progression of radiation-induced fibrosis.

As radiation-induced fibrosis is a potential side-effect of cancer radiotherapy, Nexinhib20 could only be considered therapeutically if it does not interfere with the cancer treatment. To test this, established EO771 breast cancer metastasis in the lung, where treated with fractionated radiotherapy targeted to the right lung, with Nexinhib20 or its Veh administered one day prior to the treatment and following each radiation dose (Fig. 5E). To assess treatment efficiency, metastatic burden was compared between the targeted right lung side (IR) and the non-targeted left lung side (offIR), with a ratio of one indicating no effect. While Veh-treated mice showed variable responses, all Nexinhib20-treated mice exhibited a decrease in metastatic burden in the irradiated lung (Fig. 5F). This effect was specific to radiotherapy, as the Nexinhib20 treatment itself had no impact on tumour burden in the off-target lung (Fig. 5G). These results suggest that Nexinhib20 does not impair radiotherapy efficacy and may even enhance it by producing a more consistent treatment response. Collectively, these observations support Nexinhib20 as a potential strategy to delay radiation-induced fibrosis without compromising cancer therapy.

## Discussion

Recent evidence has reshaped the traditional view of neutrophils as short-lived inflammatory cells, highlighting their broader roles in tissue repair (19). However, sustained or excessive neutrophil activation can promote fibrosis development (17). For instance, neutrophils have been implicated in renal fibrosis following sterile kidney injury (36) and studies about bleomycin-induced pulmonary fibrosis have yielded mixed results (37). While neutrophil depletion did not alter collagen deposition after bleomycin treatment (38,39), inhibition of NE reduced collagen accumulation (40,41), supporting a pro-fibrotic neutrophil phenotype. Despite these insights, the role of neutrophils in the context of radiation-induced fibrosis remains poorly defined, particularly at later stages when clinical fibrosis becomes apparent, as most studies have focused on short-term responses.

Radiotherapy can induce sterile tissue injury in non-targeted healthy regions, triggering inflammatory cascades and fibrotic tissue development. Building on previous work from our lab demonstrating that radiation exposure to healthy lung recruits neutrophils capable of inducing early perturbations in both epithelial and mesenchymal cells (29), we hypothesised that neutrophils actively contribute to the long-term development of radiation-induced pulmonary fibrosis.

To address this, we used the small animal radiation research platform (SARRP) to deliver a precise and image-guided radiation dose targeted to the lung, while minimising off-target effects and enabling long-term follow-up without clinically limiting side effects, such as skin inflammation commonly observed with thoracic-targeted irradiation. Four months after radiation exposure, fibrotic patches, characterised by excessive collagen deposition, and an accumulation of lipid-laden macrophages were observed. These foamy macrophages are known to release pro-fibrotic mediators, with their accumulation likely reflecting fibrotic progression (10–12,14). Interestingly, neutropenic mice exhibited a mitigated fibrosis phenotype, highlighting a central role of neutrophils in driving fibrosis development. Their influence is evident early, as neutrophil depletion prevents the ECM remodelling that typically occurs two weeks post-irradiation.

We previously showed that, at seven days post-radiation, neutrophils in pre-irradiated lung display increased nuclear hypersegmentation and granule protein content (29). Our findings extend these observations, showing that these features persist for at least two weeks post-irradiation. This is accompanied by the expression of markers of activation and cellular aging at the time point where we also observed measurable effects on ECM remodelling. Together, these characteristics define a radiation-educated neutrophil phenotype that contributes to fibrotic development.

Interestingly, although MPO and NE levels were increased in radiation-educated neutrophils, genetic ablation of either enzyme alone did not prevent early radiation-induced matrix-remodelling. This is in contradiction to previous studies where sivelestat-mediated NE inhibition reduced early collagen deposition in the context of the bleomycin lung fibrosis model (40,41) . Interestingly, inhibition of neutrophil degranulation with Nexinhib20 significantly reduced early ECM remodelling and fibrosis progression. Nexinhib20 treatment also seems to modulate the radiation-educated neutrophil phenotype. While their elevated activation levels appeared unaffected, the proportion of aged neutrophils, which are capable of homing back to the bone marrow (18), showed a downward trend. Furthermore, the characteristic hypersegmentation and increased motility of radiation-educated neutrophils were either absent or noticeable reduced. Interestingly, Nexinhib20 induced a strong pro-angiogenic signature in radiation-educated neutrophils while maintaining their ECM remodelling characteristics. These transcriptional changes were accompanied by a shift in the mesenchymal cell population from a pro-fibrotic to a pro-inflammatory state, favouring tissue integrity over excessive collagen deposition (Fig. 5H). Remarkably, this reprogrammed phenotype persisted after treatment withdrawal four weeks post-irradiation with fibrosis levels being strongly reduced 16 weeks after radiation exposure.

While previous studies have shown that neutrophils can deposit collagen (36,42) or transport pre-existing ECM components to the site of injury (43), our data indicate that, despite an increased ECM-related signature, neutrophils might also promote radiation-induced fibrosis through their influence on fibroblasts. The importance of neutrophil–fibroblast crosstalk in driving lung fibrosis has been demonstrated in prior work, with ARG1-mediated ornithine metabolism identified as a key profibrotic mechanism (44) . Additional evidence suggests that neutrophils can also promote fibroblast differentiation into myofibroblasts via extracellular trap formation (45) or NE release (46) .

Mechanistically, Nexinhib20 blocks the Rab27a–JFC1 interaction, preventing azurophilic granule release without disrupting core neutrophil functions such as phagocytosis or neutrophil extracellular traps formation (34). However, it remains unclear whether the observed phenotypic shift of radiation-educated neutrophils under Nexinhib20 treatment arises from direct interference with their degranulation function or as a feedback response from reduced matrix remodelling. Since the anti-fibrotic effect of neutrophils persists after cessation of the Nexinhib20 treatment, despite their short lifespan, feedback from the ECM and fibroblasts may reinforce neutrophil reprogramming toward a pro-angiogenic phenotype, thereby limiting fibrosis progression.

Importantly, Nexinhib20 treatment did not compromise the effectiveness of radiotherapy, underscoring its translational potential. Overall, this study identifies neutrophils as important drivers of radiation-induced fibrosis and highlights inhibition of neutrophil degranulation via Nexinhib20 as a therapeutic strategy to delay clinical fibrosis progression.

## Methods

### Statistics

Statistical analyses were performed in GraphPad Prism v10.6.0. Graphs were generated with Prism and schematics with BioRender. Normality was assessed with the Kolmogorov-Smirnov test or for small sample sizes with the Shapiro-Wilk test. For normally distributed data, comparisons between two groups were conducted using paired or unpaired two-tailed t-tests. Equality of variances was assessed by F-test and Welch’s correction was applied when variances differed. Non-normally distributed two-group comparisons were analysed with the Mann-Whitney test. For comparisons among more than two groups, one-way ANOVA was applied to normally distributed data, while the Kruskal-Wallis test was used otherwise. The exact statistical tests used are indicated in each figure legend. Statistical significance was defined as p < 0.05.

Data were represented in three different ways: as a superplot, a paired dot plot or a bar chart. The paired dot plots and bar charts only show the average values for each biological replicate, whereas the superplot illustrates the distribution of ROIs using truncated violin plots with the average for each biological replicate overlaid as a dot, which was also used for statistical analyses. Normally distributed data are presented with mean with its standard error of mean (± SEM), and non-normally distributed data with median and its interquartile range (± IQR), as specified in the figure legends. Sample sizes were informed by prior studies with comparable experimental designs (29,47–53) and are clearly stated in the figure legend.

### Mouse strains

Female mice aged 6–12 weeks were used in all experiments, either from one of the two WT strains (FVB/n and C57BL/6) or the three genetically modified mouse models. The WT strains were obtained from The Jackson Laboratory. The GCSFko mice, originally kindly provided by the J. Huelsken laboratory, were backcrossed to FVB/n background. Ela2-Cre knock-in mice on a C57BL/6 background were obtained from the European Mouse Mutant Archive and are referred to as Ela2ko mice. MPOko mice on a C57BL/6 background were a gift from the V. Papayannopoulos laboratory.

All mouse lines used in this study were bred and maintained under specific pathogen-free conditions at the Francis Crick Biological Research Facility. No more than five mice were housed per cage in individually ventilated cages containing wood chip bedding and nestlets, with a 12-hour light/dark cycle at 21 ± 2C and 55% ± 10% humidity. Food and water were provided ad libitum. Breeding and all animal procedures were performed in accordance with the Animals (Scientific Procedures) Act 1986 and the Animal Welfare Act 2006 at the Francis Crick Biological Research Facility, following UK Home Office regulations under project licence P83B37B3C and PP5920580.

### Cell lines

The murine breast cancer cell line EO771 was provided by the Cell Service Unit of the Francis Crick Institute, where it was routinely authenticated and tested for mycoplasma. Cells were cultured in Dulbecco’s modified eagle medium (DMEM) with 10% FBS, 20mM HEPES and 100U/mL penicillin-streptomycin at 37°C and with 5% CO_2_. Freshly thawed EO771 cells were only used for *in vivo* experiments after at least three passages.

### Experimental lung metastases models

Breast cancer metastases in the lung were generated by intravenous injection of 0.5 Mio EO771 cells into syngeneic C57BL/6 mice. Animals were monitored daily in accordance with the project licence guidelines as well as the NCRI Guidelines for the Welfare and Use of Animals in Cancer Research UK. Lungs were harvested 16 days post-injection. Mice were euthanised by cervical dislocation and lung tissue was fixed for histology. Flow cytometry of the middle lobe was performed to verify the efficiency of neutrophil depletion.

### Neutrophil depletion

In FVB/n mice, neutrophils were depleted via intraperitoneal (IP) injection of 50 μg rat anti-mouse Ly6G (αLy6G) antibody in 100 μL phosphate buffered saline (PBS) (54). Treatments began one day before radiation, followed by injections immediately after each radiation session, which were thereafter administered every other day until the experimental endpoint. Rat IgG at the same dose served as an isotype control. This antibody-based neutrophil depletion method was applied for experiments lasting up to two weeks, whereas neutropenic GCSFko mice were used for longer-term studies due to the limitations of sustained antibody-mediated depletion.

### NT degranulation inhibition with Nexinhib20

Nexinhib20 (Nex), a Rab27a-JFC1 binding inhibitor, was used to block azurophilic granule degranulation (34). Mice received IP injections at 0.03 mg/g body weight (maximum 500 μL), with Nexinhib20 dissolved in 5% Dimethyl sulfoxide (DMSO) and 12.5% Koliphor EL in PBS. Control animals received Veh injections containing the same DMSO and Koliphor EL concentrations. Injections followed the same schedule as the neutrophil depletion antibody, starting one day before the first radiation session, followed by an injection immediately after irradiation, and continuing every other day until the experimental endpoint.

### Lung-targeted X-ray irradiation (partial or whole lung)

Fibrosis-related experiments and cancer radiotherapy studies involved lung-targeted irradiation, either partial or whole lung. Irradiation was performed using the SARRP, allowing precise, CT-guided targeting. Mice were anaesthetised with isoflurane and underwent a computed tomography scan (60 kV, 0.8 mA) for treatment planning in the Muriplan software. Lung contours for the left and right lung side were defined separately to compare the radiation dose received by the two sides via the dose volume histogram. This enables accurate and consistent radiation delivery.

For partial-lung irradiation, a single isocentre was placed in the right lung lobe and a variable collimator was used to precisely fit the radiation field. Prescribed doses of 6 Gy or 13 Gy (220 kV, 13 mA, 0.15 mm Cu) were delivered via two 10 arcs with the aim that at least 90% of the targeted right lung tissue received approximately 5 Gy or 11 Gy, respectively, while 10% of the non-targeted left lung tissue was exposed to 1 Gy or less. Whole-lung irradiation followed the same approach, with a separate isocentre being positioned in each lung side. Mice remained anaesthetised on a heated bed throughout imaging and radiation delivery. Non-irradiated mice were anaesthetised for same amount of time, approximately 15-20min per mouse.

### Lung tissue digestion

For MACS of neutrophils or fibroblasts, the harvested lung tissue was minced and enzymatically digested in 1 mL Hanks’ Balanced Salt Solution (HBSS) containing liberase Tm [0.25 U/mL], liberase Th [0.25 U/mL], and 1% DNase I [5 μL/mL] at 37C with continuous shaking at 180 rpm for 30 min. For flow cytometry, minced lung tissue was passed directly through a 100 μm strainer to generate a cell suspension, which was centrifuged at 300 × g for 7 minutes. Red blood cells were removed using 1:10 diluted RBC lysis buffer for 5 minutes at room temperature. The resulting pellet was washed with MACS buffer (PBS containing 0.5% bovine serum albumin and 2.5 mM EDTA) and filtered through a 35 μm strainer into a 5 mL FACS tube to obtain a single-cell suspension for staining and analysis.

### Flow cytometry analysis

Prior to staining, single-cell suspensions from lung tissue were incubated with mouse FcR Blocking Reagent (1:20 in MACS buffer) for 10 minutes at room temperature to prevent non-specific binding. Cells were then labelled with fluorescent-conjugated antibodies (Table 3) at a 1:100 dilution in MACS buffer on ice for 20 minutes, protected from light. Cell viability was assessed using either 4’,6-diamidino-2-phenylindole (DAPI) (1:500 in MACS buffer) or Zombie Aqua (1:300 in PBS), following two washes with the respective buffer. After viability staining, cells were washed once with MACS buffer and resuspended in 200 μL of fresh MACS buffer for acquisition. Samples were analysed on a BD LSR-Fortessa cytometer, and data were processed using FlowJo v.10.4.2. Median fluorescence intensity (MFI) was used to quantify surface marker expression. The gating strategy for neutrophil identification in lung is shown in Supplementary Information Fig. 1.

### Magnetic Activated Cell sorting (MACS)

Neutrophils were isolated from previously digested lung tissue using MACS positive selection for Ly6G. Each lung was processed individually, with cells first incubated for 5 minutes at room temperature in 80 μL FcR-blocking reagent (1:20 dilution in MACS buffer). Ly6G MicroBeads (20 μL per sample) were then added, mixed by pipetting, and incubated on ice for 15 minutes. For live imaging assays, the αLy6G-AF647 antibody was administered 5 minutes after MicroBead application. Samples were washed twice with MACS buffer and resuspended in 500 μL MACS buffer before loading onto pre-wetted LS columns for magnetic separation, as per the manufacturer’s instructions. Isolated neutrophils were counted and viability assessed prior to downstream assays.

For fibroblast isolation, the Ly6G-negative flow-through was stained with CD45, EpCAM and CD31 MicroBeads followed by a procedure analogous to the Ly6G MACS workflow. Fibroblasts collected in the flow-through while unwanted cells remained bound to the column.

### Sample preparation for bulk RNAseq

For bulk RNA sequencing, neutrophil and fibroblast populations were collected from non-irradiated control lungs and from irradiated lungs treated with either Veh or Nexinhib20 four weeks post-irradiation, as described above. RNA was extracted using the Direct-zol RNA MicroPrep kit from zymo following the manufacturer’s instructions. Libraries were prepared with the Watchmaker RNA Ribo/Globin Depletion kit, and sequencing was performed on a NovaSeq X platform targeting 25 million reads per sample (50 bp paired-end sequencing read length).

### Neutrophil speed assay on gelatin-coated matrix

To compare the motility of radiation-educated versus control neutrophils, αLy6G-AF647- stained cells were plated on top of Oregon Green™ 488-conjugated gelatine matrix. For ECM preparation, 24-wel plates (Cellvis P241.5HN) were treated with 100% Nitric acid for 10min.

Following three PBS washes, the wells were coated with 0.005% Poly-L-Lysine in PBS (100 µL per well) at room temperature. After 20 minutes, the solution was removed and prewarmed Oregon Green™ 488 gelatin (1 mg/mL, 50 µL) was applied to cover each well for 20 minutes in the dark. Wells were treated with ice-cold 0.5% glutaraldehyde (50 µL) for 15 minutes on ice and an additional 15 minutes at room temperature, washed thrice with PBS and quenched with 20 mM glycine for 5 minutes. After further PBS washes, wells were incubated with 1 mL 70% ethanol for 5 minutes, washed again and equilibrated with fresh media for 1 hour. Wells were maintained in PBS at 37C overnight before adding 50–60 µL of neutrophil suspension (8,000 cells per well) in neutrophil media (phenol-red-free DMEM/F12 supplemented with B- 27, 20 ng/mL EGF, 20 ng/mL FGF and 4 μg/mL heparin). Neutrophils were allowed to settle for 4 hrs and then imaged every 20 min for 7h using a Nikon ECLIPSE Ti2 microscope (20x magnification). For analysis, IMARIS (v10.2.0) was used. Briefly, the 647nm channel was used to detect spots of an estimated diameter of 6μm and quality above 1.9400. tracking was performed using the Autoregressive Motion algorithm. Generated tracks were inspected for accuracy and if necessary, manually rearranged using the Edit Tracks tool. Tracks that lasted for less than 3 frames (1h) were excluded from analysis.

### Neutrophil immunofluorescence staining

To assess neutrophil nuclear morphology and MPO levels after radiation or Nexinhib20 treatment, 30,000 MACS-isolated neutrophils in 50 µL PBS were seeded onto poly-L-lysine– coated coverslips. After allowing cells to settle at room temperature for 20min, they were fixed with 200 µL of 4% paraformaldehyde for 10 minutes. Following three washes with PBS, coverslips were treated with a blocking solution containing 1% bovine serum albumin, 10% FCS, and 0.2% Triton X-100 for 30 minutes, followed by a 10-minute incubation with mouse FcR Blocking Reagent (1:20 in PBS) to prevent non-specific antibody binding. For MPO detection, primary antibody (1:200 in blocking solution) was applied overnight at 4C. The next day, coverslips were washed three times with PBS before incubation with the corresponding secondary antibody (anti-goat-AF555, 1:200) and DAPI (1:500) in blocking buffer for 45 minutes at room temperature. After final washes and a brief water rinse, coverslips were mounted onto slides, air-dried and stored at 4C until imaging. Z-stack images were acquired using an Olympus CSU-W1 SoRa Spinning Disk confocal microscope with a UPLAPO OHR 60x/1.5 oil-immersion objective. Images were deconvoluted using 5D Adaptive Deconvolution Maximum Likelihood Estimation and projected into 2D sum images for each channel in FIJI.

Analysis was conducted in QuPath. MPO content per neutrophil was quantified from the 2D sum images. Each neutrophil area was defined by its MPO signal and mean fluorescence intensity (MFI) of MPO-AF555 per cell was calculated. Cells in close proximity or at the image border were excluded. To correct for background and enable comparisons between images, three 117.36 μm² ROIs were drawn per image and their average MPO-MFI was subtracted from the corresponding neutrophil-MFI.

Hypersegmented neutrophils were identified using the Z-stack images to examine nuclear morphology in 3D. Neutrophils with six or more nuclear lobes were classified as hypersegmented. Cells that were difficult to identify were noted but not counted (∼5% of neutrophils/image).

### Lung tissue staining

For histological quantification, lung tissue was fixed overnight in 10% neutral buffered formalin. It was processed and embedded in paraffin. Prior to paraffin embedding, tissue samples underwent automated processing, which comprised dehydration, clearing and infiltration with embedding medium. Sections of 4 μm thickness were cut at intervals of ∼150 μm. Before staining, slides were incubated at 60C for 1 hour, then deparaffinised and rehydrated using an automated stainer.

Hematoxylin and eosin (H&E) staining was performed with an automated Tissue-Tek Prisma Stainer. Slides were imaged on a Zeiss Axio Scan.Z1, and tumour burden per section was quantified using StrataQuest software with a pre-trained semi-automated protocol.

For collagen visualisation, slides were placed in picrosirius red solution for 60 minutes, followed by differentiation in 0.5% acetic acid for 5 minutes. Slides were dehydrated through graded ethanol (95% and 100%), cleared in xylene and mounted. Imaging was performed on the Zeiss Axio Scan.Z1 and fibrotic areas and foamy macrophage coverage were quantified either manually in an unbiased manner or by using the FIJI TWOMBLI plugin.

### Immunohistochemistry staining for neutrophils

Immunohistochemistry staining was done to detect neutrophils in lung tissue. For this purpose, sections underwent antigen retrieval in a solution containing 0.1% trypsin and 0.1% CaCl₂ in 1X Tris-buffered saline (pH 7.6) at 37C for 15 min. Background peroxidase activity was quenched by treating the sections with 1.6% hydrogen peroxide (H₂O₂) in PBS for 10 min before rinsing in water. Prior to primary antibody staining, lung tissue sections were placed in the blocking solution for 30 minutes at room temperature. To stain for the neutrophils, S100a9 (1:1000 in blocking solution) was applied overnight at 4C. After three washes on the following day, secondary antibody staining with a goat anti-rat-Bio (diluted 1:275 in blocking solution) was done for 45min at room temperature. Signal amplification was achieved using the ABC (Avidin-Biotin Complex) kit and visualisation of the antigen-antibody complexes was achieved by the 3,3’-Diaminobenzidine (DAB) substrate. Sections were counterstained for tissue contrast and coverslipped using an automated stainer. Images were captured on a Zeiss Axio Scan.Z1, and DAB-positive neutrophils were quantified using QuPath.

### Immunofluorescence staining of lung sections

For immunofluorescence staining of the foamy macrophages, slides underwent heat-mediated antigen retrieval in 10 mM Tris-EDTA buffer (pH 9). The buffer was pre-heated to 90C for 8 min before adding the slides, which were then maintained at 45C for 15 min, after which the solution cooled gradually. Once at room temperature, the blocking solution was applied for 30min, followed by overnight incubation with primary antibodies diluted in blocking solution at 4C. On the next day, slides were washed and incubated for 2 h at room temperature with fluorophore-conjugated secondary antibodies (1:200) and DAPI (1:500) in blocking solution. After further washing, Sudan Black (0.1% in 70% ethanol) was applied for 20 min, then rinsed off before mounting. Imaging was performed on a Zeiss Upright710 confocal or Axio Scan.Z1 microscope, and quantification was done with QuPath.

### Image analysis

Most image analyses were performed in QuPath and FIJI as outlined above, including quantification of neutrophils in IHC and IF-stained sections. Tumour burden on H&E slides was assessed with StrataQuest.

To assess the matrix changes due to irradiation, picrosirius red stained lung sections were used. Fibrotic areas and regions occupied by foamy macrophages were manually quantified in an unbiased manner over three levels per sample, while the ECM areas were evaluated using the TWOMBLI plugin in FIJI (9). For this purpose, 30 ROIs were generated for the IR and offIR lung samples, with equal distribution of the IR-ROIs between the cranial, middle, and caudal lobes. The accessory lobe was left out, as its position can vary towards the mediastinum and does not reliably fall within the irradiated field. The selected ROIs were analysed with the FIJI macro TWOMBLI (9), which combines several FIJI plugins. Ridge Detection (55) was used for unbiased identification of filamentous structures, while Anamorf (56) and OrientationJ (57) provided further quantification of the detected matrix. The tool outputs the reconstructed matrix and associated parameters, such as matrix area, which reflects the amount of matrix per ROI and accounts for fibre thickness. This measure was used as the main readout in this study. Analyses were performed at one level per biological replicate. Data are displayed as a superplot with violin plots showing the distribution of all ROIs per condition and the mean value for each mouse indicated as a dot. For paired analysis, averages per biological replicate were plotted, with lines linking the off-IR (left) and IR (right) lung sides of the same animal.

### Bulk RNAseq analysis

Bulk RNAseq data from isolated lung neutrophils and the mesenchymal cell population were processed using the nf-core/rnaseq pipeline (version 3.18.0) with Nextflow (version 24.10.2) and Singularity (version 3.11.3). Reads were aligned to the mus musculus reference genome (GRCm38, downloaded from Ensembl) using the RSEM-STAR option within the pipeline to generate gene-level count matrices. R (version 4.5.1, RStudio version 2025.05.1+513) was used for gene count import into DESeq2 (version 1.49.2) for differential gene expression analysis, after pre-filtering genes with fewer than ten total counts across. Heatmaps of differentially expressed genes were generated using the pheatmap R package (version 1.0.13) based on variance-stabilising transformation–normalised expression values.

For gene set enrichment analysis, genes were considered significant if the baseMean ≥ 5 and the adjusted p-value ≤ 0.05. Log₂ fold-change thresholds of ≥ 0.5 for mesenchymal cells and ≥ 1 for neutrophils were applied. Functional enrichment analysis was performed using the STRING database (version 12.0) (35), focusing on Biological Process Gene Ontology (GO) terms. Enriched terms were ranked by enrichment score (signal), and the top 10 were visualised. GO terms sharing a similarity score ≥ 0.5 were clustered, and together with their false discovery rate (FDR) values and gene counts, are illustrated in the enrichment visualisation figure.

## Acknowledgements

We thank Adam Karoutas, Xuanxuan Fan, Rute M.M. Ferreira, Nicolas Rabas, Felipe S. Rodrigues and Stefania Di Blasio for their support, helpful discussions and technical assistance throughout this project. We are grateful to the core facilities at the Francis Crick Institute that assisted this work, including the Biological Research Facility (Adebambo Adekoya, Hannah Easter, Scott Lighterness, Daiva Poniskaitiene, Matthew William Knowles, Antonio Summa, Veronika Smirnova), the In Vivo Imaging facility (Thomas Snoeks, Peter Johnson), the Experimental Histopathology Laboratory (Richard Stone and Emma Nye), the Bioinformatics Facility (James Campbell, Probir Chakravarty), the Advanced Sequencing Facility, the Flow Cytometry Facility, the Advanced Light Microscopy Facility and the Cell Service Unit. Schematics were created with BioRender. This work was supported by the Francis Crick Institute, which receives its core funding from Cancer Research UK (CC2051), the UK Medical Research Council (CC2051) and the Wellcome Trust (CC2051). Funding was also received from the European Research Council grant (ERC CoG-H2020725492). Parts of this study, including some experimental data and methodological details, are adapted from the author’s doctoral thesis (Stefanie Ruhland, 2025, King’s College London).

## Authors contribution

S.R designed and performed the experiments, analysed and interpreted the data and wrote the manuscript. K.N. performed and analysed the neutrophil motility experiment and provided valuable discussions. V.L.B. provided experimental and technical support, managed colony breeding and provided valuable discussions. T.R. provided experimental support. E.S. provided support with TWOMBLI analysis. J.B. provided technical support with the SARRP.

E.N. and A.E.A. participated in the design of the study and provided valuable discussions. I.M. designed, supervised and funded the study, interpreted the data and edited the manuscript. K.N., V.L.B and A.E.A reviewed the manuscript and gave valuable feedback.

