## Supplementary Figures for "Inhibition of neutrophil degranulation by Nexinhib20 delays the development of radiation-induced pulmonary fibrosis"

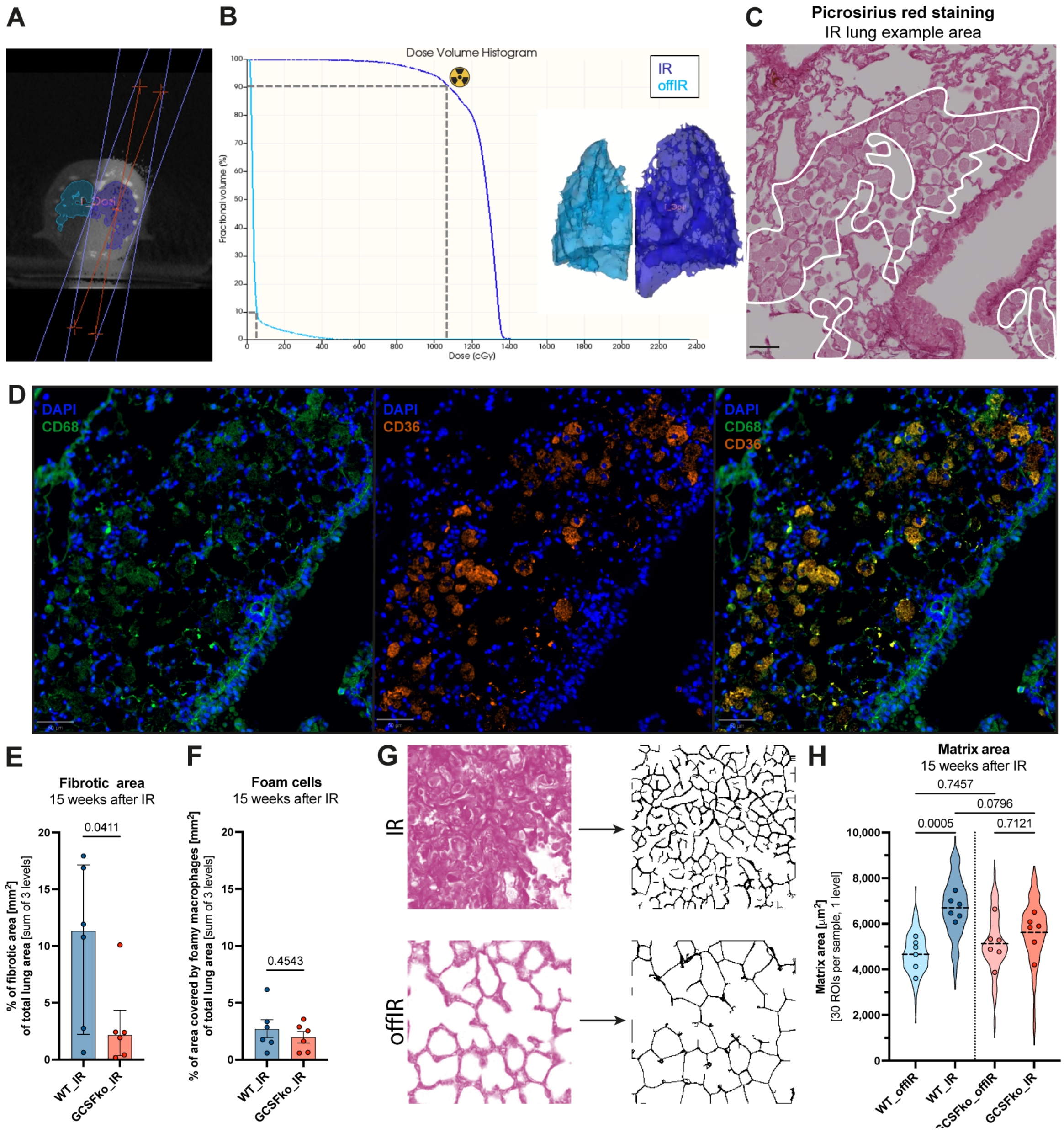


**Supplementary Figure 1: Neutrophils support the development of clinical radiation-induced fibrosis. (A)** Representative CT scan of the setup for targeted irradiation of the right lung using the SARRP. **(B)** Dose volume histogram from a representative mouse illustrating the proportion of lung tissue receiving specific radiation doses. The setup aims for ~90% of the targeted right lung (IR) to receive approximately 11 Gy, while minimizing off-target exposure in the left lung (offIR), where 10% of tissue receives ≤1 Gy. **(C)** Representative image of picrosirius red stained lung tissue from the irradiated right lung, showing areas with foam cell detection. Scale bar: 50 µm. **(D)** Immunofluorescence of the same region as in (C). The first image shows DAPI (blue) and CD68 (green), the second shows DAPI with CD36 (red) and the third is an overlay of all three channels. Scale bar: 50 µm. **(E–F)** Data from Fig. 1E with bar charts showing individual mouse data for the sum of fibrotic area (E) and foamy macrophage-covered area (F) in the IR lung over three levels. Statistical analysis was performed using the Mann–Whitney test (E) and unpaired t-test (F). Data are presented with median ± IQR (E) or mean ± SEM (F) (n = 6). **(G)** Representative images of ROIs and their corresponding matrix generated using TWOMBLI. The top panel shows an ROI from IR lung tissue with matrix accumulation, while the bottom panel shows an ROI from non-targeted (offIR) lung tissue. **(H)** Quantification of matrix area across 30 ROIs (48,163 µm²) from one level per sample using TWOMBLI. Violin plot shows the distribution of ROIs and dots indicate the average value for each mouse. Statistical analysis was performed on individual mice using one-way ANOVA. Data are presented with mean (n = 6).

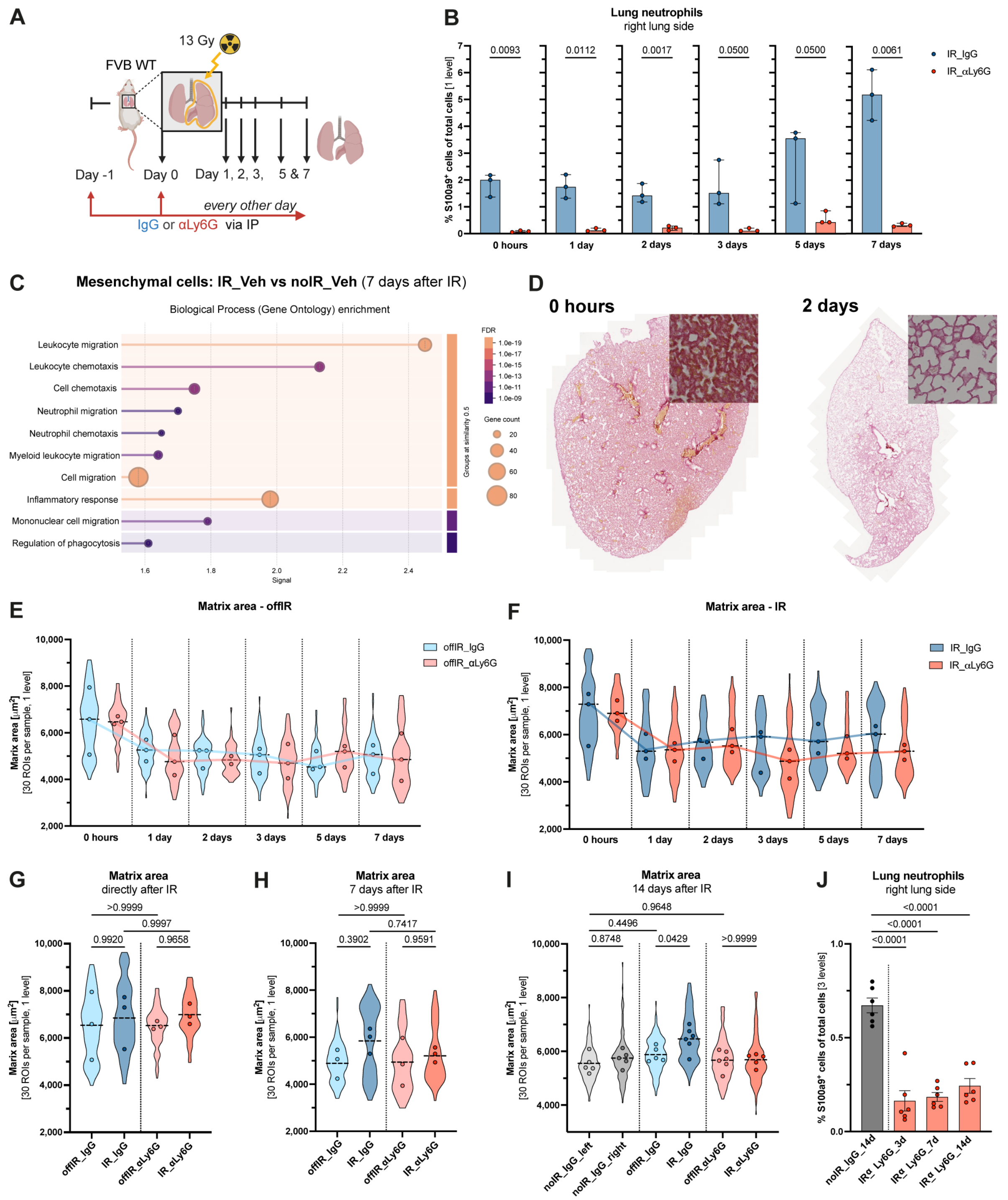


**Supplementary Figure 2: Neutrophils support the development of radiation-induced early ECM changes. (A)** Schematic illustrating experimental set-up to evaluate the role of neutrophils on early matrix changes after radiation exposure. **(B)** Quantification of neutrophil presence in lung tissue by counting S100a9⁺ cells as a percentage of total cells from a single level of the IR lung tissue. Tissue was collected immediately, 1, 2, 3, 5 or 7 days after radiation exposure. Statistical analysis was performed using Welch's t-test, Welch's t-test, unpaired t-test, Mann–Whitney test, Mann–Whitney test and Welch's t-test, respectively. Data are presented with median ± IQR (n = 3). **(C)** GO-term analysis of differential expressed gene analysis of bulk RNAseq of mesenchymal cells isolated from IR vs noIR lung tissue seven days post-irradiation via the STRING pathway analysis webpage (1). Data was generated by Nolan et al. (2022) (2) and reanalysed. **(D)** Representative image of picrosirius red stained IR lung tissue collected either immediately after irradiation or two days afterwards. **(E and F)** Quantification of matrix area across 30 ROIs (48,163 µm²) from one level per sample for offIR lung (E) and IR lung side (F) using TWOMBLI. Violin plots show the distribution of ROIs and dots indicate the average value for each mouse. Data are presented with median, with a line indicating changes in the median over time (n = 3). (G and H) Data from E and F comparing offIR and IR for lungs collected immediately after **(G)** or seven days after radiation exposure **(H).** Statistical analysis was performed using one-way ANOVA. Data are presented with mean (n = 3). **(I)** Day 14 data from Fig. 1G, with added comparison to offIR lung tissue. Statistical analysis was performed using one-way ANOVA. Data are presented with mean (n = 5-6). (J) Quantification of neutrophil presence in lung tissue by counting S100a9$^+$ stained cells as a percentage of total cells across slides from three levels. Statistical analysis was performed using one-way ANOVA. Data are presented with mean ± SEM (n = 5-6).

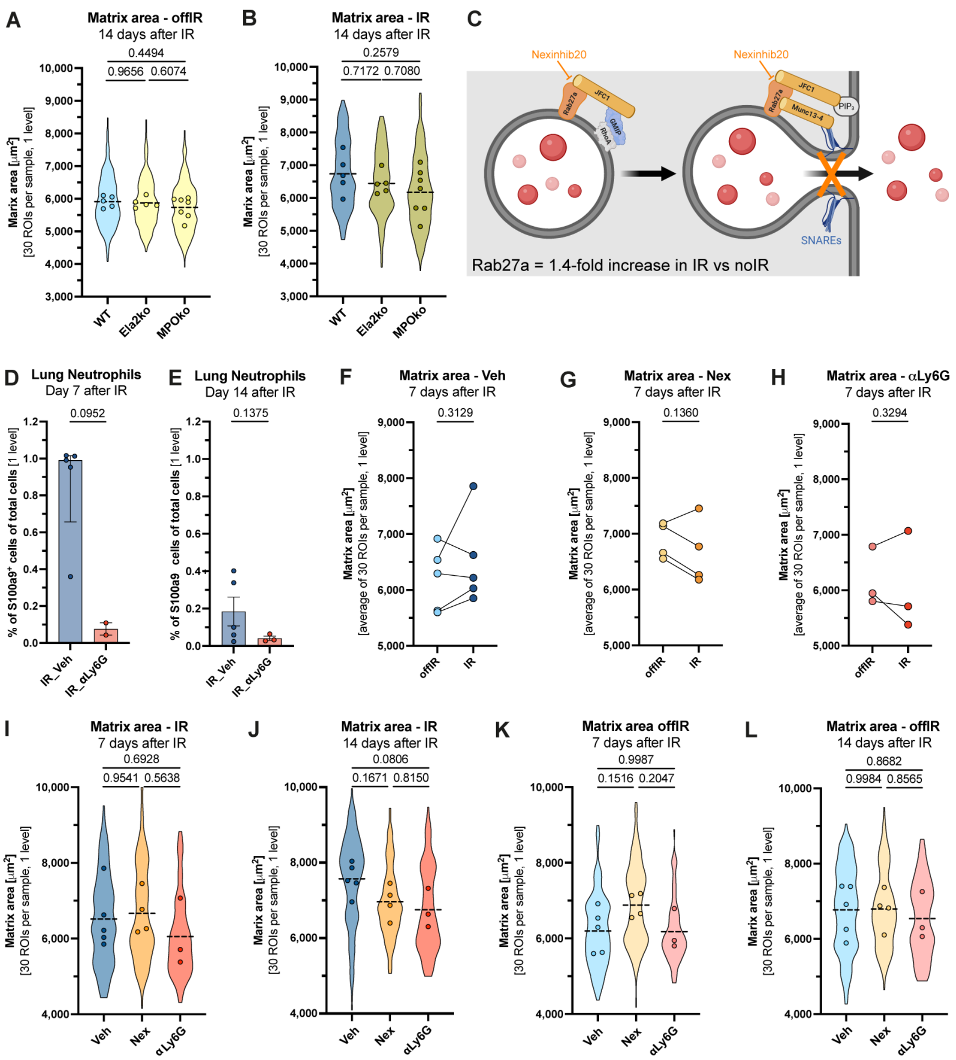


**Supplementary Figure 3: Blocking degranulation with Nexinhib20 prevents early matrix changes. (A and B)** Data from Fig. 3B-D with quantification of matrix area in offIR (A) or IR (B) lung tissue 14 days post-irradiation across 30 ROIs (48,163 µm²) from one level per sample using TWOMBLI. Violin plot shows the distribution of ROIs and dots indicate the average value for each mouse. Statistical analysis was performed using one-way ANOVA. Data are presented with mean (n = 5-6). **(C)** Schematic illustrating the effect of the degranulation inhibitor Nexinhib20, which block the binding of Rab27a and JFC1 and thus prevents the degranulation of azurophilic granules (Schematic was adapted (3–5) ). Rab27a was 1.4-fold upregulated in radiation-educated neutrophils compared to controls (reanalysis of neutrophil proteomics data from Nolan et. al (2022) (2) ). **(D and E)** Quantification of neutrophil presence in lung tissue by counting S100a9⁺ stained cells as a percentage of total cells across slides from one level either 7 (D) or 14 days after irradiation (E). Statistical analysis was performed using Mann-Whitney test (D) or Welch's t test (E). Data are presented with median ± IQR (D) or mean ± SEM (E) (n = 3-5). **(F-H)** Quantification of matrix area across 30 ROIs (48,163 µm²) from one level per sample using TWOMBLI. Dots show the average per mouse, with paired offIR and IR lungs indicated. Statistical analysis was performed using paired t-test (n = 3-5). **(I-L)** Data from Fig. 3F-H and Supp. Fig. 3F-H with quantification of matrix area in IR (I and J) or offIR (K and L) lung tissue either 7 (I and K) or 14 days post-irradiation (J and L) across 30 ROIs (48,163 µm²) from one level per sample using TWOMBLI. Violin plot shows the distribution of ROIs and dots indicate the average value for each mouse. Statistical analysis was performed using one-way ANOVA. Data are presented with mean (n = 3-5).

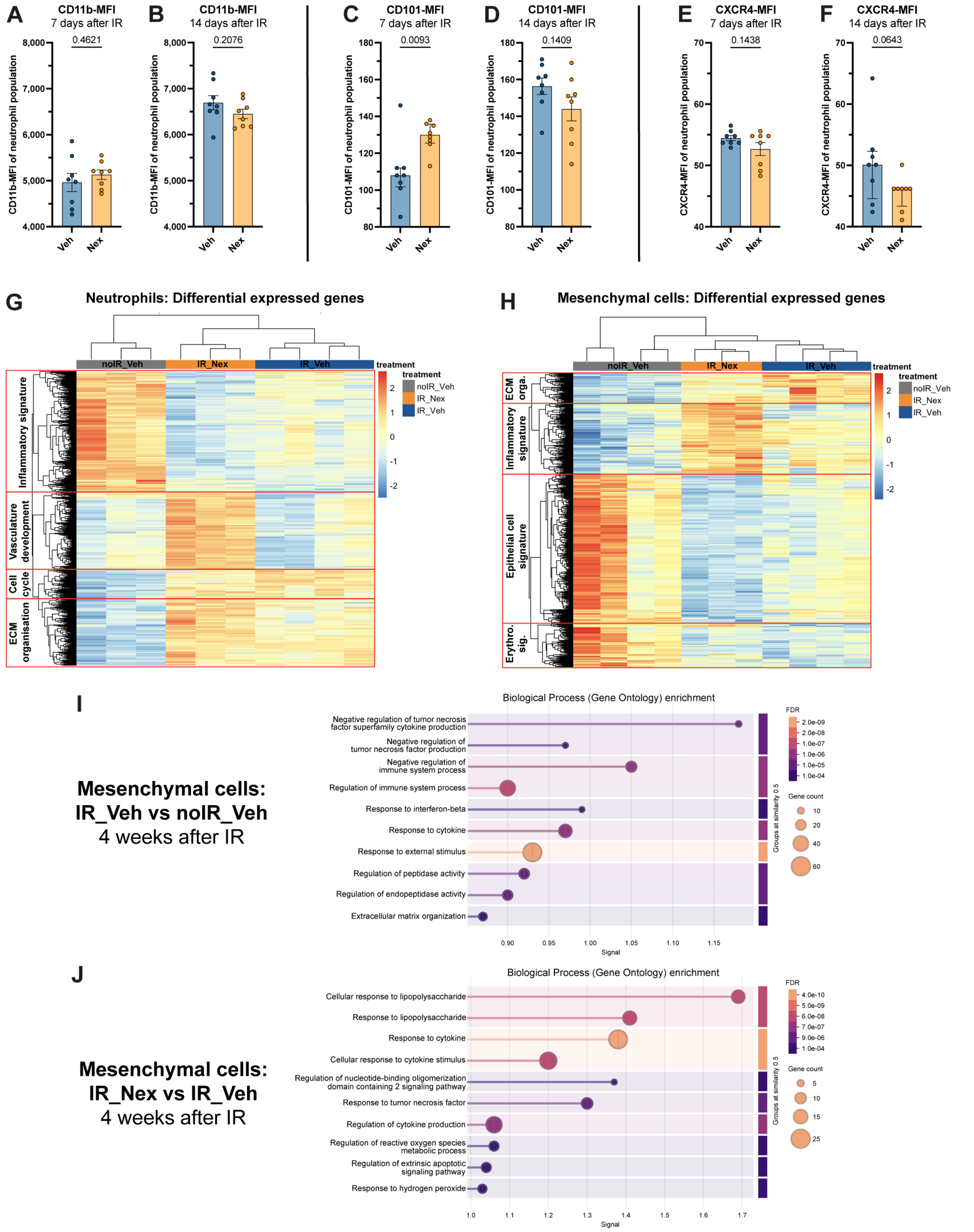


**Supplementary Figure 4: Nexinhib20 modulates the radiation-educated neutrophil phenotype and transcriptomic signature. (A-F)** Expression levels neutrophil surface markers CD11b (A and B), CD101 (C and D) and CXCR4 (E and F) quantified by FACS. Neutrophils were either isolated from pre-irradiated lung tissue either 7 (A, C and E) or 14 days (B, D and F) after radiation exposure. Statistical analysis was performed using unpaired t-test (A, B, D), Mann-Whitney test (C and F) or Welsh's t-test (E). Data are presented with mean ± SEM (A, B, D and E) or median ± IQR (C and F). **(G and H)** Heatmap showing the expression of differential expressed genes among the neutrophil (G) or mesenchymal population (H) isolated either from noIR lung samples, Veh- or Nex-treated IR lung samples. **(I and J)** GO-term analysis of differential expressed gene analysis of bulk RNAseq of mesenchymal cells isolated from either Veh-treated (I) or Nex-treated IR (J) vs Veh-treated noIR lung tissue four weeks post-irradiation via the STRING pathway analysis webpage (1).

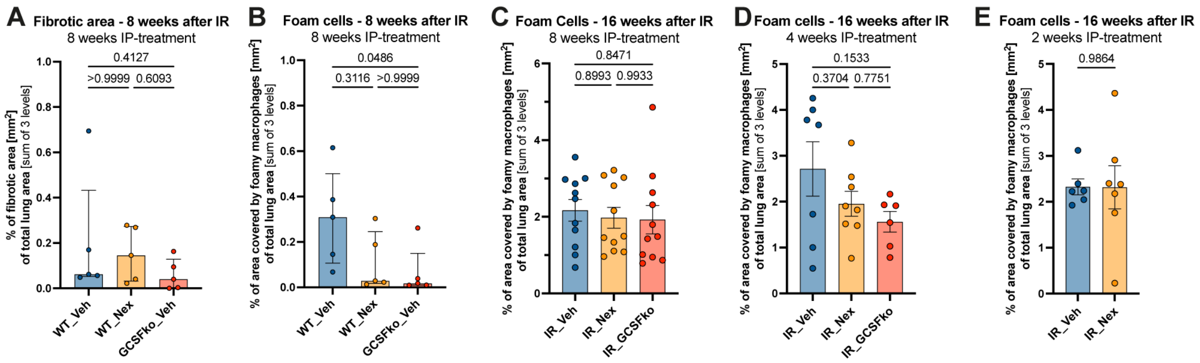


**Supplementary Figure 5: Nexinhib20 as a potential therapeutic strategy to delay radiation-induced fibrosis without compromising cancer radiotherapy efficacy. (A-E)** Bar charts showing individual mouse data for the sum of fibrotic area (A) and foamy macrophage-covered area (B-E) in the IR lung over three levels. Mice received Nexinhib20 or Vehicle for 8 weeks and lungs were harvested either 8 weeks (A and B) or 16 weeks post-irradiation (C-E). Statistical analysis was performed using Kruskal-Wallis test (A and B), one-way ANOVA (C and D) or Welsh's t-test (E). Data are presented with median ± IQR (n = 5) for A and B and with mean ± SEM (n = 6-11) for C-E.
