## Supplementary material for "Inhibition of neutrophil degranulation by Nexinhib20 delays the development of radiation-induced pulmonary fibrosis": Materials

### ***Key resource table***

| REAGENT | SOURCE | IDENTIFIER |
|---|---|---|
| **Antibodies (IF and IHC)** | | |
| Goat anti-humane/mouse-MPO (1:200) | R&D | AF3667 |
| Rabbit anti-mouse CD68 (1:100) [EPR23917-164] | Abcam | ab283654 |
| Mouse anti-mouse CD36 [JC63.1] | Abcam | ab23680 |
| Rat anti-mouse S100a9 (1:1000) | In house | N/A |
| Donkey anti-mouse-AF555 (1:200) | Thermo Fisher Scientific | A-31570 |
| Donkey anti-rabbit-AF488 (1:200) | Thermo Fisher Scientific | A-212-6 |
| Goat anti-rat-IgG (1:275) | Thermo Fisher Scientific | 31471 |
| anti-mouse-AF555 | Thermo Fisher Scientific | A-31570 |
| **Antibodies (Flow Cytometry, MACS)** | | |
| CD45-BV421 [30-F11] | BioLegend | 103134 |
| CD11b-FITC [M1/70] | BioLegend | 101206 |
| CD101-PE [Moushi101] | Thermo Fisher Scientific | 12-1011-82 |
| CD184 (CXCR4)-BV421 [2B11] | BD Horizon | 562738 |
| Ly6G-BUV395 [1A8] | BD Horizon | 563978 |
| Ly6G-AF647 [1A8] | BioLegend | 127610 |
| Ly6G MicroBeads | Miltenyi Biotec | 130-120-337 |
| **In vivo treatment** | | |
| InVivoPlus anti-mouse Ly6G [1A8] | BioXCell | BP0075-1 |
| IgG | in house | Y13/238 |
| Nexinhib20 | Tocris | 6089 |
| **Chemicals** | | |
| Kolliphor EL | Sigma-Aldrich | C5135-500G |
| DMSO | Sigma-Aldrich | D2650-100ML |
| 4',6-diamidino-2-phenylindole (DAPI) | Sigma-Aldrich | D9542 |
| Liberase TH | Roche | 5401127001 |
| Liberase TM | Roche | 5401151001 |
| Deoxyribonuclease I (DNAse I) | Merck Sigma-Aldrich | DN25-100MG |
| Zombie NIR | BioLegend | 423106 |
| 0.4% trypan blue solution | Thermo Fisher Scientific | 15250061 |
| EDTA | Sigma-Aldrich | ED-100G |
| Red Blood Cell Lysis Buffer | Miltenyi Biotec | 130-094-183 |
| FcR-blocking reagent | Miltenyi Biotec | 130-092-575 |
| TritonX-100 | Sigma-Aldrich | X100-1L |
| 10% NBF | Sigma-Aldrich | HT501128-4L |
| Pierce$^{TM}$16 % Formaldehyde | Sigma Aldrich | 28908 |
| VFM Harris Haematoxylin Stain (Acidified) | CellPath | RBA-4205-00A |

| | | |
|---|---|---|
| Eosin Y Solution | CellPath | RBC-0100-00A |
| Scott's Tap Water Substitute | In house | N/A |
| Xylene | Fisher Chemical | X/0200/17 |
| Weigert's Iron Hematoxylin | Sigma-Aldrich | HT1079 |
| Biebrich Scarlet-Acid Fuchsin Solution | Sigma-Aldrich | HT151-250 |
| Ethanol | Sigma-Aldrich | 32205-2.5L-M |
| Aniline Blue Solution | Sigma-Aldrich | HT154-250 |
| Picro Sirius Red Stain Kit | Abcam | ab150681 |
| Acetic acid | Sigma Aldrich | A6283-100ML |
| Tris EDTA | TaKaRa Bio | T9131 |
| Hydrogen peroxide solution ($H_2O_2$) | Merck | H1009-5ML |
| Trypsin | Sigma Aldrich | T4799-5G |
| Vectastain Elite ABC Kit | Vector Laboratories | PK-6100 |
| DAB Peroxidase Substrate Kit | Vector Laboratories | SK-4100 |
| Gelatin From Pig Skin, Oregon Green 488 Conjugate | Thermo Fischer Scientific | G13186 |
| Glutaraldehyde | Sigma-Aldrich | 340855 |
| Glycine | In house | / |
| HCS CellMask™ Stain – 555nm | Thermo Fischer Scientific | H32713 |
| **Reagents for media** | | |
| FBS | Labtech | FB-1001/100-500 |
| HBSS | Thermo Fisher Scientific | 14025092 |
| DMEM | Thermo Fisher Scientific | 41965062 |
| DMEM Phenol red free | Thermo Fisher Scientific | 11580846 |
| DMEM/F12 | Thermo Fisher Scientific | 12634-010 |
| Human EGF | Thermo Fisher Scientific | PHG0313 |
| Human FGF-10 | Thermo Fisher Scientific | 100-26-100UG |
| Heparin | Sigma-Aldrich | 9041-08-1 |
| B-27 Supplement | Thermo Fisher Scientific | 17504044 |
| GlutaMAX$^{TM}$ Supplement | Thermo Fisher Scientific | 35050061 |
| Trypsin-EDTA (0.25%), phenol red | Thermo Fisher Scientific | 25200056 |
| Penicillin-Streptomycin (5,000 U/mL) | Thermo Fisher Scientific | 11528876 |
| Insulin solution human | Sigma-Aldrich | I9278 |
| HEPES (1M) | Thermo Fisher Scientific | 15630080 |
| Bovine Serum Albumin (BSA) | Sigma-Aldrich | A7906-500G |
| **Mouse strains** | | |
| C57BL/6 | The Jackson Laboratory, supplied by The Francis Crick Institute | N/A |
| FVB/n | The Jackson Laboratory, supplied by The Francis Crick Institute | Jax Strain # 001800 |
| GCSFko FVB/n | gift from the J. Huelsken laboratory and backcrossed to FVB/n | N/A |
| Ela2ko | the European Mouse Mutant Archive | N/A |

| | | |
|---|---|---|
| MPOko | gift from the V. Papayannopoulos laboratory | N/A |
| **Other supplies** | | |
| Poly-L-lysine coated coverslip | Corning | 354085 |

| **Software and algorithms** | | |
|---|---|---|
| BioRender | BioRender | https://www.biorender.com |
| Fiji/ImageJ v. 2.14.0/1.54f | Open source (Schindelin et al., 2012) | https://imagej.nih.gov/ij/ |
| FlowJo software v.10.4.2 | BD Life Science | https://www.flowjo.com |
| GraphPad Prism v.10.3.1 | GraphPad | https://www.graphpad.com |
| Muriplan v.3.0.1 | Xstrahl | https://xstrahl.github.io/muriplan/ |
| QuPath v. 0.4.2. | Open source (Bankhead et al., 2017) | https://qupath.github.io/ |
| R v. 4.4.0. | Open source | https://cran.r-project.org/ |
| RStudio v. 2024.04.1+748 | Open source | https://posit.co/products/open-source/rstudio/ |
| StrataQuest | TissueGnotics | https://tissuegnostics.com/products/contextual-imageanalysis/strataquest |
