## Supplementary Information for "Inhibition of neutrophil degranulation by Nexinhib20 delays the development of radiation-induced pulmonary fibrosis"

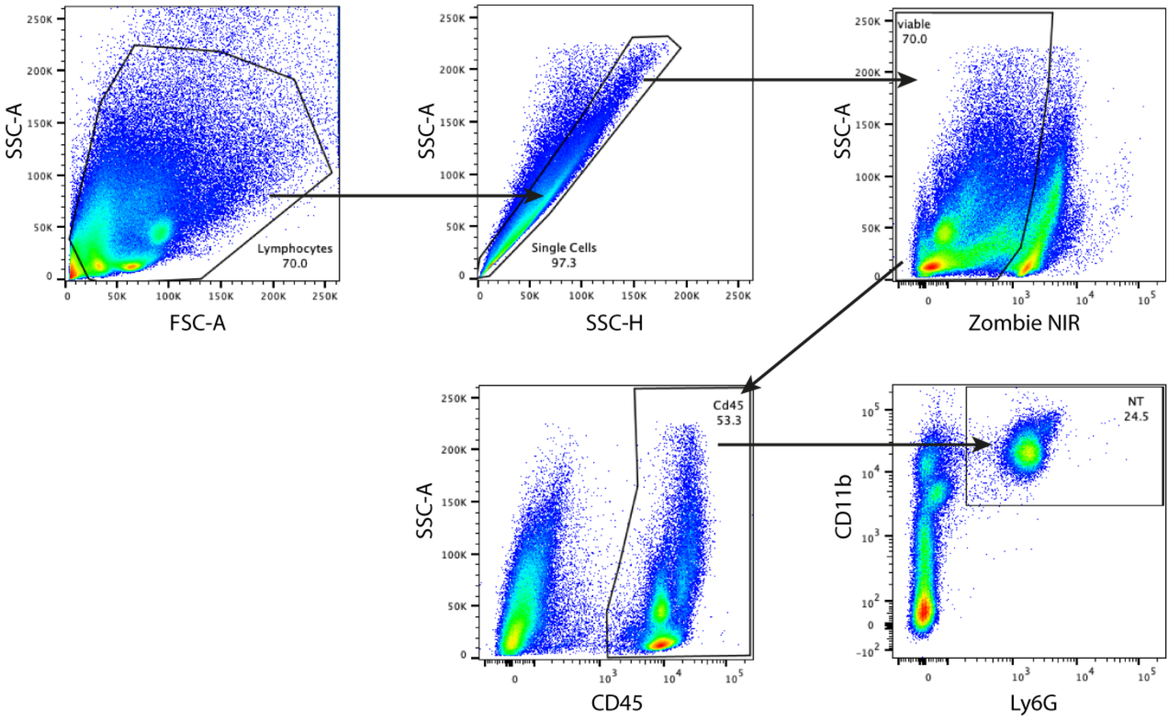


**Figure 1: FACS gating strategy** Example of gating strategy for neutrophil identification. First, we gated on lymphocytes, then selected for single cells by SSC-A vs SSC-H. Next, we gated on viable cells, identified by using Zombie NIR, followed by gating on $CD45^+$ immune cells and finally identification of $CD11b^+$ and $Ly6G^+$ neutrophils.
